# Relaxation of cardiac pericytes by GLP-1 activating K_ATP_ channels mediates remote ischaemic preconditioning cardioprotection

**DOI:** 10.1101/2025.06.26.661857

**Authors:** Svetlana Mastitskaya, Felipe Santos Simões de Freitas, Lowri E. Evans, David Attwell

## Abstract

Failure to reperfuse the coronary microvasculature (“no-reflow”) affects up to 50% of patients after unblocking a coronary artery that caused ischaemia and acute myocardial infarction. No-reflow is associated with reduced left ventricular ejection fraction, increased infarct size and death. We have established that no-reflow results from cardiac pericytes constricting coronary capillaries, and that pharmacologically relaxing pericytes reduces no-reflow. Remote ischaemic preconditioning, by briefly making a limb ischaemic, protects against cardiac ischaemic injury, and we have shown this is mediated by release of the gut hormone glucagon-like peptide 1 (GLP-1). We now demonstrate that, by releasing GLP-1, remote ischaemic preconditioning reduces pericyte-mediated coronary capillary constriction and no-reflow, and that the dilating effect of GLP-1 on coronary capillaries is abolished by block or genetic deletion of pericyte K_ATP_ channels. These results define a brain-gut-heart pathway mediating remote ischaemic cardioprotection, and suggest pharmacological therapies to reduce ischaemia-induced coronary no-reflow and improve post-infarct recovery.

## Introduction

Partial or complete occlusion of a coronary artery, often following atherosclerotic plaque rupture, leads to cardiac ischaemia with ST segment elevation on the ECG. Primary percutaneous coronary intervention (PPCI) is the preferred treatment and aims to restore blood flow in the affected artery and minimize damage to the heart muscle. However, this does not guarantee reperfusion of the downstream capillaries supplying the myocardium^1,2^ and a lack of capillary reperfusion - ‘no-reflow’ - affects up to 50% of patients^3,4^.

Infarct size (the amount of damaged heart tissue) is a predictor of adverse events and left ventricular remodeling after a heart attack. Cardioprotection studies frequently focus on reducing infarct size, but recent research suggests that the presence of no-reflow is an independent predictor of adverse outcomes and may be more significant than infarct size itself^4^. However, while many studies have examined infarct size, only a few have specifically addressed the microvascular occlusion causing no-reflow as a therapeutic target^5,6^. This is despite a meta-analysis^7^ demonstrating a robust relationship between microvessel occlusion and mortality or hospitalisation for subsequent heart failure within one year, with a 1% increase in vessel occlusion predicting 14% and 11% increases in death and hospitalisation respectively. Understanding and targeting therapeutically the mechanism(s) leading to vessel occlusion may therefore significantly improve outcome after cardiac ischaemia.

We have previously shown that pericyte-mediated capillary constriction contributes significantly to no-reflow after cardiac ischaemia^8^. For the brain, where the same mechanism operates, this results from a rise of intracellular calcium concentration, [Ca^2+^], triggering pericyte contraction during ischaemia, with subsequent pericyte dysfunction or death maintaining the constriction when the arterial blood supply is restored^9^. Similar events occur following renal ischaemia^10^. Despite the critical role of pericyte-mediated capillary constriction not yet being broadly known in the cardiac community^11^, this finding has been independently reproduced for the heart^12^. We have shown that pericyte-mediated no-reflow can be significantly reduced by infusing adenosine at the time of reperfusion^8^, and it has also been suggested that remote ischaemic pre-conditioning (RPc, in which a limb is made transiently ischaemic before the cardiac ischaemia) has a cardioprotective action that is mediated by relaxation of pericytes^12^.

These results suggest that, if the mechanism by which RPc relaxes coronary pericytes could be defined, it might offer a pharmacological therapeutic approach superior to the use of adenosine which lowers blood pressure and has other side effects. We have previously discovered that the cardioprotective action of RPc is mediated by a soluble factor released by activity in the motor fibres of the posterior gastric branch of the vagus nerve^13,14^, which was suggested^15^ to be glucagon-like peptide 1 (GLP-1). By examining cardiac ischaemia *in vivo* and imaging pericyte-mediated coronary capillary constriction in the pressurised vasculature of excised ventricle, we now demonstrate that the effect of RPc on coronary blood flow involves release of GLP-1, which relaxes coronary pericytes by activating K_ATP_ channels in these cells. The magnitude of the resulting blood flow increase and cardiac protection can be modulated by agents that alter trafficking of K_ATP_ channels to the surface membrane, including AMP kinase, NO and acetylcholine.

## Results

### GLP-1 receptors are on pericytes on coronary capillaries

The fluorescent GLP-1R-binding compound^16^ LUXendin555 bound to pericytes (and also some endothelial cells) in the mouse coronary vasculature, suggesting the presence of GLP-1 receptors on pericytes (Fig. 1a). Similarly, an antibody against GLP-1R also labelled mouse and cultured human coronary pericytes (Fig. 1b-c).

**Figure 1:**
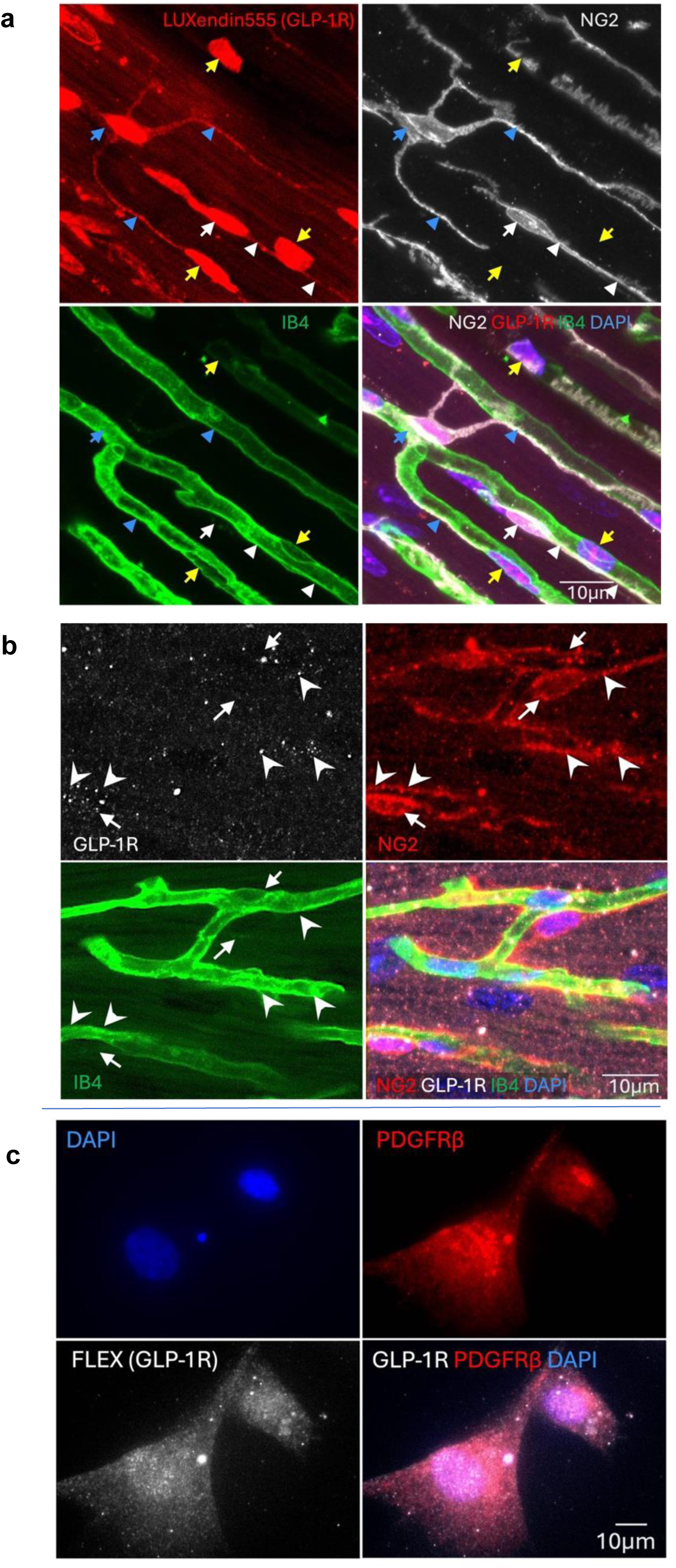
Cardiac pericytes express GLP-1 receptors. (**a**) Labelling of mouse coronary microvasculature with LUXendin555 (red) which labels GLP-1R. NG2 antibody labelling (white) shows that LUXendin labelled cells are pericytes (white arrow and arrowheads show somata and processes respectively), telocytes (blue arrow/arrowheads, which may function similarly to pericytes but have processes contacting more than one capillary) or endothelial cells (yellow arrows). IB4 labelling green) shows the basement membrane around endothelial cells and pericytes; The merge of these labels (bottom right) also includes nuclear labelling with DAPI (blue). (**b**) Labelling of mouse coronary microvasculature with anti-GLP-1R antibody (white), NG2 (red), IB4 (green) and DAPI (blue). White arrows and arrowheads show pericyte somata and processes respectively. (**c**) Labelling of cultured human cardiac pericytes with Fluorescein-Trp25-Exendin-4 (FLEX) probe binding to GLP-1R, along with PDGFRβ (red) and nuclear (DAPI, blue) labelling.

### No-reflow after coronary ischaemia reflects capillary constriction

Occluding the left anterior descending coronary artery (LAD) for 45 mins and then allowing 15 mins reperfusion induced a long-lasting 40% decrease of coronary blood volume in the part of the left ventricle supplied by the LAD, as compared with sham-operated rats (p=0.0031, Fig. 2a, c). This correlated with an increase in the percentage of capillaries blocked in the core of the area at risk (1-2 mm deep in the left ventricular wall: Fig. 2b, f) from 11.6±1.5% (n=3 hearts) to 73.9±9.0% (n=6, significantly different, p=0.0001, Tukey multiple comparisons test).

**Figure 2.**
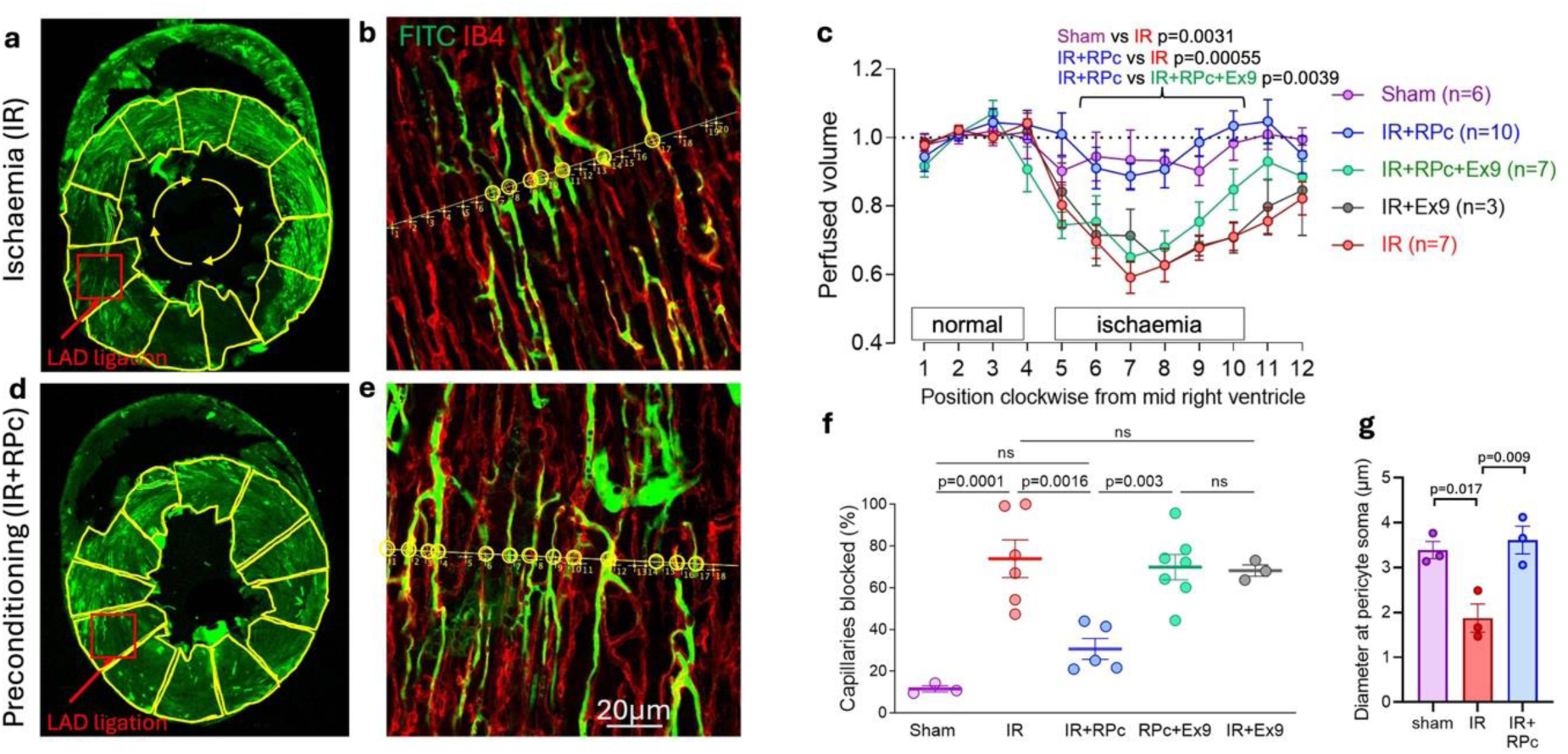
RPc prevents ischaemia/reperfusion induced capillary constriction and no-reflow, via GLP-1R activation. (**a, b**) Low power (a) and zoomed in view (b) of area outlined red in a) of a cross-section of heart after ischaemia/reperfusion (top). In (b), a line is drawn perpendicular to the capillaries, and the ratio of number of blocked to total number of capillaries crossed by the line is calculated. Perfused capillaries are marked by circles. (**c**) Perfusion volume assessed as mean intensity of FITC-albumin fluorescence (green) in each region of interest (ROI) outlined in yellow in (a). ROIs are indexed with numbers starting in the middle of the interventricular septum and proceeding clockwise around the left ventricular wall (as seen from above); intensities are normalized to the average intensity of the first 3 ROIs. (**d, e**) As in (a-b) but with RPc applied before ischaemia, which restores perfusion volume (c). (**f**) Percentage of capillaries blocked in the anterior wall of the left ventricle is increased by ischaemia/reperfusion, but this increase is reduced by RPc. The inhibition of capillary block and rescue of flow (c) by RPc is blocked by the selective GLP-1R blocker Exendin(9-39) (Ex9). Ex9 had no significant effect in ischaemia unless RPc was performed. Data are mean ± s.e.m. (**g**) Diameter of capillaries at pericyte somata after sham ischaemia, after ischaemia (IR), and after ischaemia preceded by remote ischaemic preconditioning (IR+RPc). P values compare sham and IR+RPc conditions with ischaemia alone (IR). P values are corrected for multiple comparisons.

We have previously shown that this capillary block and reduction of perfused blood volume are produced by pericyte-mediated constriction^8^. As previously, the diameter of capillaries at pericyte somata was greatly reduced (by ∼45%) after ischaemia and reperfusion (p=0.017 by one-way Anova in 48 capillaries in 3 sham-operated hearts and 48 capillaries in 3 hearts exposed to ischaemia-reperfusion, Fig. 2g).

### RPc reduces no-reflow via GLP-1R mediated pericyte relaxation

Remote ischaemic preconditioning prevented the ischaemia-evoked pericyte-mediated capillary constriction (p=0.009, Fig. 2g), and (presumably as a result) reduced capillary blockage from 73.9±9.0% to 30.7±5.0% of capillaries blocked (p=0.0016, in 5 hearts) compared to coronary ischaemia/reperfusion alone (Fig. 2d-f). RPc also restored the post-ischaemic perfused blood volume (p=0.00055 compared with ischaemia/reperfusion alone) to a level similar to that seen for sham-operated mice (Fig. 2c).

Applying the GLP-1R blocker Exendin(9-39) (Ex9) intravenously, before the ischaemia, prevented RPc-mediated reduction of capillary block (resulting in 69.9±6.1% capillary block, n=7 hearts, p=0.003 compared with RPc+ischaemia/reperfusion, Fig. 2f), and led to an ischaemia-evoked fall of perfusion volume similar to that seen with ischaemia/reperfusion alone (p=0.0039 compared to RPc+Ex9 group, Fig. 2c). If applied with ischaemia/reperfusion in the absence of RPc, Ex9 had no effect (Fig. 2c, f), consistent with it acting solely by preventing the effects of RPc.

### GLP-1R evoked pericyte relaxation is via activation of K_ATP_ channels

To investigate the mechanism by which GLP-1 receptor activation protects against the ischaemia/reperfusion induced pericyte contraction^8,12^ that blocks capillaries and reduces coronary blood flow, we used an *ex vivo* preparation. This comprised mouse free right ventricular wall, which we perfused through the vasculature by cannulating the right coronary artery (diameter ∼150 μm, see Methods and Fig. 3a) in addition to superfusing it with extracellular fluid. Oxygen and glucose deprivation (OGD) for 25 mins evoked pericyte contraction and a constriction of capillaries, with a decrease of capillary diameter by 15.9±1.7% (p=0.0002, at n=21 pericytes in tissue from 6 hearts, compared to the initial diameter, Fig. 3b). In continuous OGD conditions, the capillary diameter continued to gradually decline to 23.0±2.6% less than the initial value (Fig. 3b).

**Figure 3.**
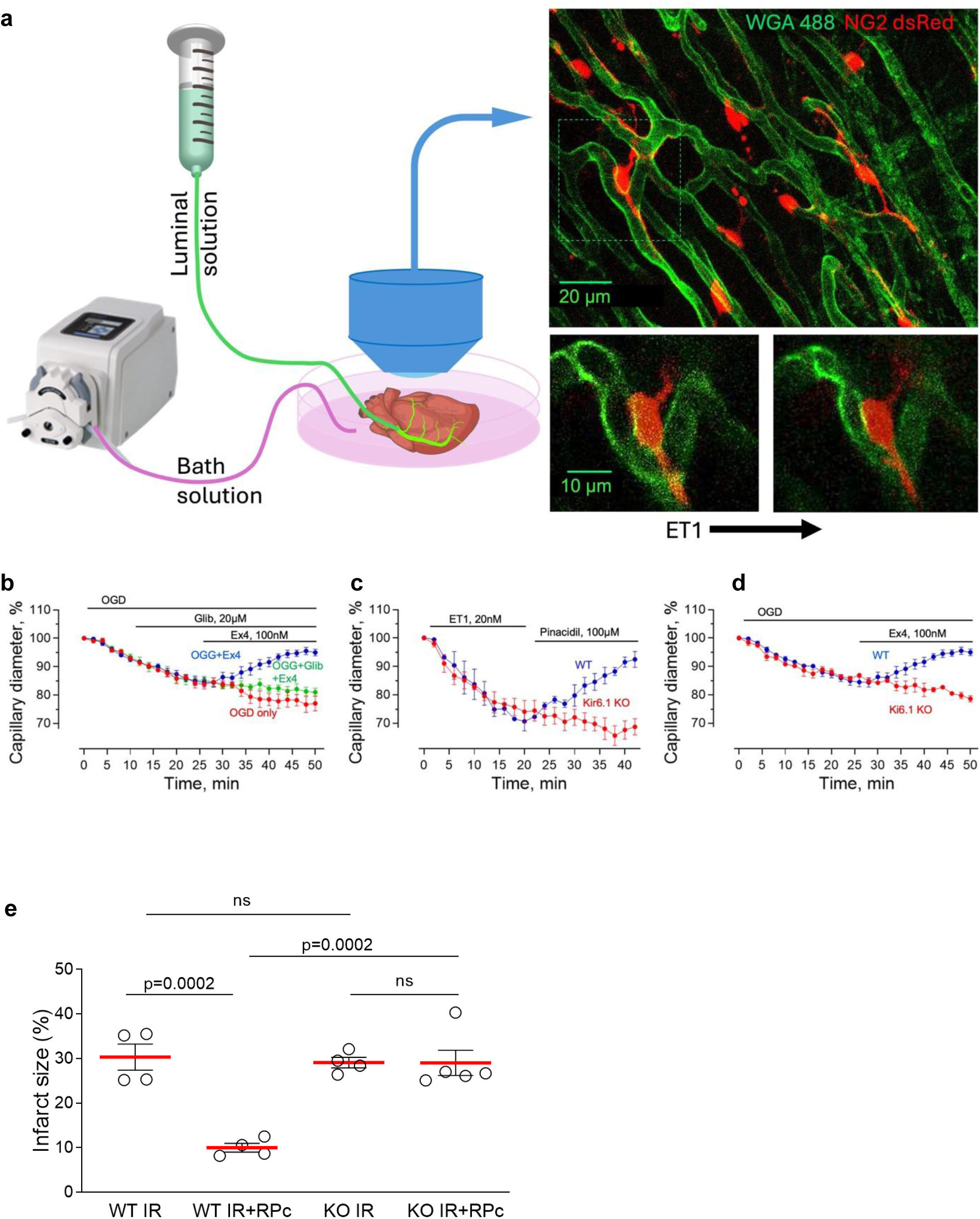
GLP-1 receptors dilate capillaries by activating K_ATP_ channels. (**a**) Schematic of the perfused right ventricle section used for these experiments: a piece of right ventricular wall is pinned to the perfusion chamber, and the right coronary artery is cannulated and perfused with a luminal solution while the tissue is superfused with a bath solution. The capillary diameters at pericyte somata are measured over time during application of vasoactive agents (e.g. endothelin-1, ET1). (**b**) Effect of oxygen-glucose deprivation (OGD, red) on capillary diameter at pericytes in ventricular tissue (21 pericytes in 6 hearts) is reversed by the GLP-1 receptor activator Exendin-4 (Ex4, 25 pericytes in 7 hearts, blue), and this rescue of diameter is greatly reduced by the K_ATP_ channel blocker glibenclamide (20 μM, 23 pericytes in 6 hearts, green). (**c**) Constricting effect of endothelin-1 (ET-1) on capillary diameter at pericytes in wild-type (WT) tissue (9 pericytes in 3 hearts) was reversed by the K_ATP_ activator pinacidil (100 μM, blue) but this reversal was abolished by knock-out of the K_ATP_ component Kir6.1 (12 pericytes in 4 hearts, red). (**d**) Constricting effect of OGD on capillary diameter at pericytes in wild-type (WT) tissue (25 pericytes in 7 hearts, blue) was reversed by Ex4, but this rescue of diameter was prevented by KO of Kir6.1 (9 pericytes in 3 hearts, red). (**e**) Myocardial infarct size evoked by 30 mins ischaemia followed by 90 mins reperfusion *in vivo* in mice (IR) was reduced by RPc, and this was abolished in mice with Kir6.1 knocked out, consistent with a role for ATP-gated K^+^ channels in mediating the effects of GLP-1 during RPc.

Activating GLP-1R with Exendin-4 (Ex4, 100 nM) in the continued presence of OGD, starting after 25 mins of OGD, relaxed pericytes within 25 mins and dilated capillaries back to a diameter that was similar to the baseline value before ischaemia/reperfusion (Fig. 3b, constricted by 5.4±1.0% at n=25 pericytes in 7 hearts, compared to 23.0±2.6% in continuous OGD without Ex4 at n=21 pericytes in 6 hearts; p=6.1×10^-6^, number of hearts was used as the statistical unit). However, in the presence of the ATP-gated K^+^ channel blocker glibenclamide (20 μM), the GLP-1R-mediated pericyte relaxation was abolished, and capillaries remained constricted by 19.0±1.4% of the initial diameter (at n=23 pericytes in 6 hearts, p=1.2×10^-4^ compared to Ex4 only, Fig. 3b). This is consistent with RPc-evoked cardiac protection being blocked by glibenclamide *in vivo*^17^ and with GLP-1 activating K_ATP_ channels in smooth muscle cells^18^ and pancreatic beta cells^19^.

ATP-sensitive K^+^ channels in vascular mural cells (smooth muscle cells around arterioles and pericytes around capillaries) have previously been shown to play a major role in regulating vascular tone^20^. Ventricular cardiac pericytes express^21^ both Kir6.1 (a key component of ATP-gated K^+^ channels^20,22^) and SUR2 (an essential subunit of coronary pericyte K_ATP_ channels^23^). To confirm the involvement of ATP-gated K^+^ channels in the regulation of capillary diameter by pericytes, pericytes were preconstricted for 20 mins with endothelin-1 (ET1, 20 nM, applied via the capillary lumen). ET1-evoked constriction is long lasting, and even after washout of the drug pericytes remained constricted for over 45 mins (data not shown). ET1 decreased the capillary diameter at pericyte somata by 29.6±3.3% (at 7 pericytes in 2 hearts after 20 mins, Fig. 3c). Applying the K_ATP_ channel opener pinacidil^22^ (100 µM, in the capillary lumen) after termination of ET1 application resulted in a relaxation of pericytes and an increase of capillary diameter back to the baseline level within 20 mins (Fig. 3c). In contrast, in tissue from mice with Kir6.1 knocked out in pericytes (see Methods), capillaries remained constricted in the presence of pinacidil (by 31.3±2.9% of the initial diameter, at n=12 pericytes in 4 hearts, compared to 7.5±2.8% in WT, p=0.007, Fig. 3c). Similarly, the reversal of OGD-induced capillary constriction by Ex4 (Fig 3b) was inhibited when Kir6.1 was knocked out in pericytes, so that capillaries remained constricted after 50 mins of OGD at 21.4±1.1% of the initial diameter, at n=9 pericytes in 3 hearts, p=1.7×10^-5^ compared to WT, Fig. 3d). Consistent with these results in isolated ventricles suggesting a role for ATP-gated K^+^ channels in mediating the effects of GLP-1 during RPc, in mice with Kir6.1 knocked out in pericytes, RPc was not protective against ischaemia/reperfusion injury (Fig. 3e).

### Modulation of the GLP-1 - K_ATP_ pathway by AMP kinase, NO and ACh

The ability of GLP-1 released by RPc to relax cardiac pericytes and increase coronary blood flow presumably depends on the activation of K_ATP_ channels in the surface membrane of the pericytes, which will hyperpolarise the pericytes and decrease voltage-gated Ca^2+^ entry. AMP-dependent protein kinase (AMPK) contributes to cell survival during cardiac ischaemia^24^ and is known to promote activation of K_ATP_ channels^25^, at least in part by increasing trafficking of these channels to the surface membrane^26^. We found that applying compound C (CC, 6-[4-(2-Piperidin-1-yl-ethoxy)-phenyl)]-3-pyridin-4-yl-pyyrazolo[1,5-a] pyrimidine), an AMPK blocker^27^ (5 µM), inhibited the GLP-1R-mediated reversal of OGD-induced pericyte-mediated constriction of coronary capillaries (p=6.4×10^-4^ at 15 pericytes in 4 hearts, Fig. 4a). This is consistent with compound C inhibiting the cardiac protection provided by remote ischaemic conditioning, resulting in a larger infarct size^28^.

**Figure 4.**
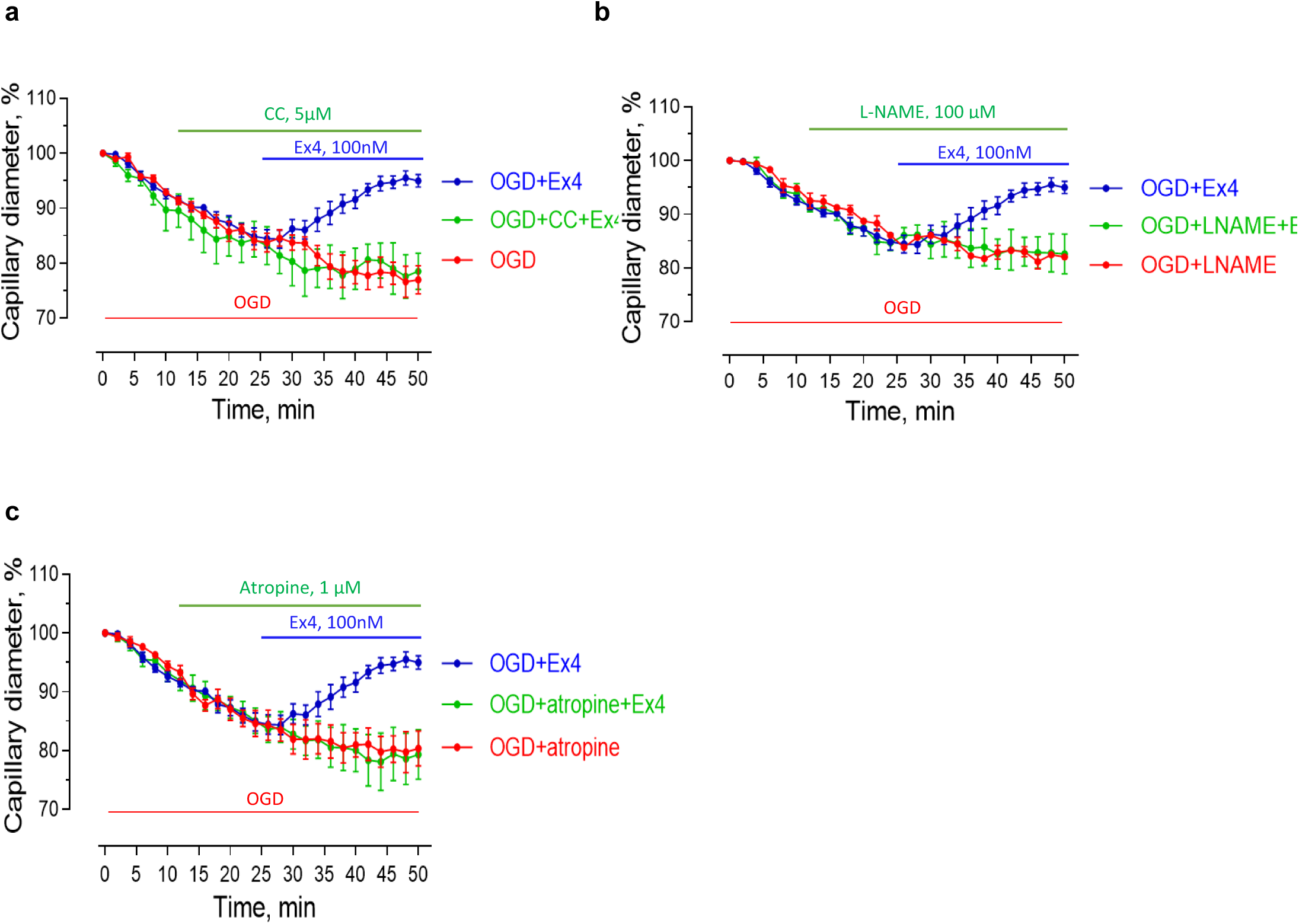
GLP-1 evoked capillary dilation in ischaemia is modulated by AMP kinase, NO and cholinergic signalling. (**a**) Reversal by Exendin-4 (25 pericytes in 7 hearts) of the OGD-evoked constriction of capillaries at pericytes (21 pericytes in 6 hearts) is abolished by the AMPK blocker compound C (CC, 5µM, 15 pericytes in 4 hearts). (**b**) The NOS blocker L-NAME (100 µM) has no effect on the OGD-evoked constriction, but abolishes the dilation evoked in OGD by GLP-1R activation with Ex4 (25, 8 and 19 pericytes in 7, 2 and 3 hearts for OGD+Ex4, OGD +L-NAME and OGD+L-NAME+Ex4, respectively). (**c**) Atropine (1 μM) has no effect on the OGD-evoked constriction, but blocks the dilation in OGD evoked by GLP-1R activation (26, 13 and 25 pericytes in 7, 4 and 5 hearts for OGD+Ex-4, OGD+atropine and OGD+atropine+Ex4, respectively).

The nitric oxide synthase (NOS) blocker L-NAME (100 µM) did not affect the constriction evoked by OGD alone, but blocked the capillary dilation evoked by activating GLP-1 receptors in OGD (p=0.0037 at 19 pericytes in 3 hearts, Fig. 4b). This is consistent with NOS block preventing the beneficial effect of RPc on no-reflow *in vivo*^29^. NO might be expected to dilate blood vessels directly by relaxing capillary pericytes or preventing them constricting^9^, but this would be expected to occur even in the absence of GLP-1 receptor activation, predicting a constriction when L-NAME is applied, which is not seen in Fig. 4b. Here, it is possible that the effect of NOS block reflects a decrease of trafficking of K_ATP_ channels to the membrane because NO has been reported to activate AMPK^30,31^. Atropine (1 μM) also did not affect the constriction evoked by OGD alone, but blocked the capillary dilation evoked by activating GLP-1 receptors in OGD (p=0.0033 at 25 pericytes in 5 hearts, Fig. 4c), consistent with atropine blocking RPc-evoked cardiac protection after ischaemia^15^. Again, the lack of effect on the OGD-evoked constriction but suppression of the Ex4 evoked dilation might reflect acetylcholine promoting trafficking of K_ATP_ channels to the membrane, because the type 3 muscarinic receptors that are cardioprotective in RPc^15^ are reported to increase activation of AMPK^32,33^.

To assess whether AMPK does indeed regulate K_ATP_ channel trafficking, we used an antibody to Kir6.1 which forms an essential subunit of coronary pericyte K_ATP_ channels^23^ (Fig. 5a). In mouse hearts subjected to 40 mins LAD ligation followed by 10 mins reperfusion, the peak fluorescence intensity of Kir6.1 labelling shifted inside the cell in animals administered with the AMPK inhibitor compound C (CC, 10 mg kg^-1^ i.p.) both in ischaemia/reperfusion (1.11±0.03 µm in the IR+CC group compared to 0.75±0.03 µm in the IR control; p=0.001), and in the animals pre-treated with Exendin-4 (Ex4, 5 µg kg^-1^ i.p.; 1.13±0.06 µm in the IR+CC+Ex4 group vs 0.67±0.07 µm in the IR+Ex4 group; p=0.0004) (Fig. 5b-c).

**Figure 5.**
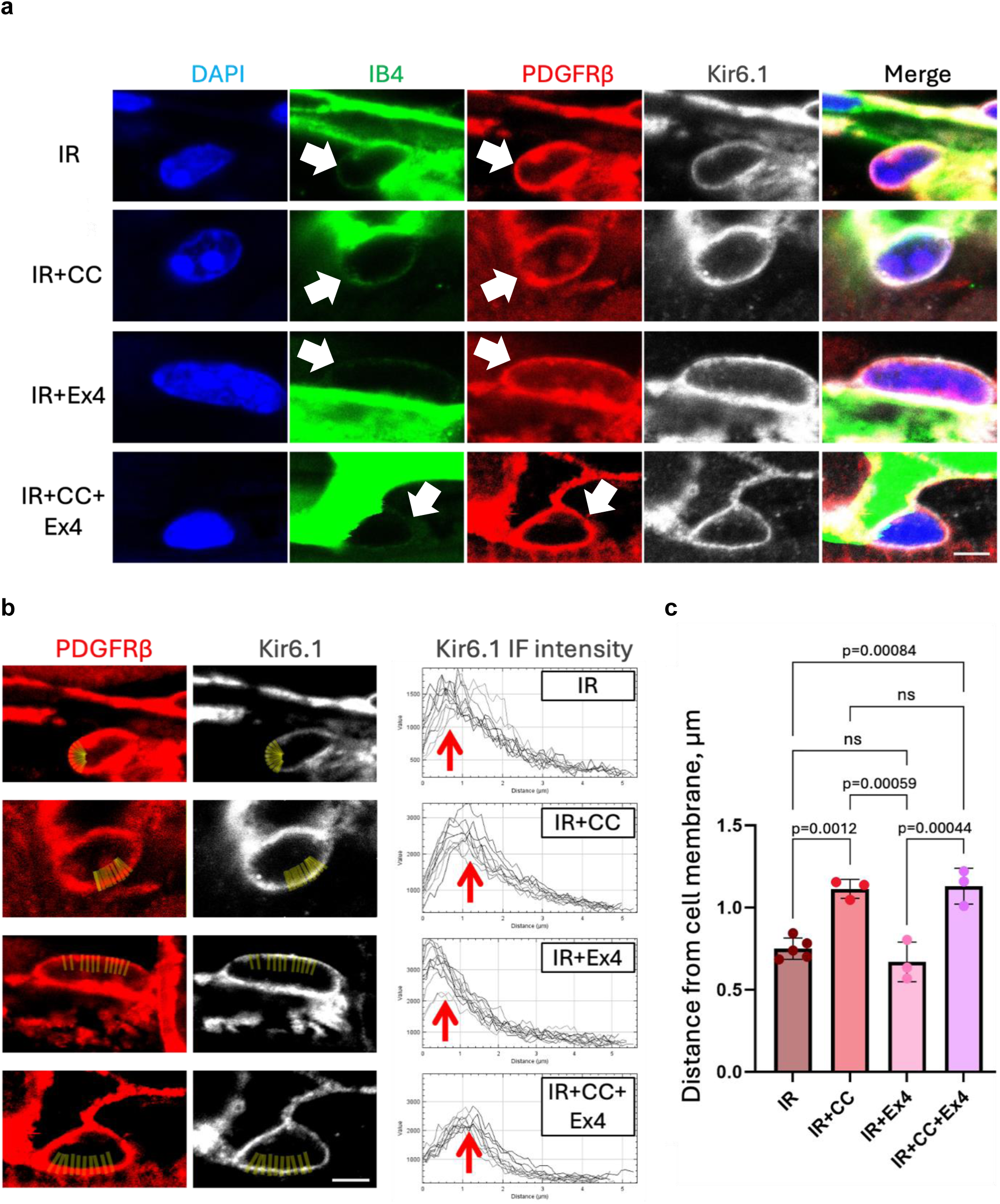
AMPK inhibition reduces trafficking of K_ATP_ channels to the cell surface. (**a**) Examples of immunolabelled cardiac pericytes from four experimental groups of mice with LAD ligation: ischaemia/reperfusion only (IR), IR group pre-treated with Compound C (IR+CC), IR group pre-treated with Exendin-4 (IR+Ex4), and IR with both CC and Ex4 (IR+CC+Ex4). IB4 labelling extends all around the pericytes (white arrows). (**b**) Examples of Kir6.1 immunofluorescence intensity profiles measured using imageJ in individual pericytes, with PDGFRβ immunostaining serving as a reference point to define the cell surface. 10 line regions of interest (ROIs, fine yellow lines) per pericyte were drawn in the PDGFRβ channel starting from the cell surface directed towards the centre of the cell, the ROIs were then transferred to the Kir6.1 channel and fluorescence intensity along each line was measured and averaged per pericyte. The distance from the membrane to the peak Kir6.1 immunofluorescence was then averaged per heart (n=40, 43, 30, 31 pericytes in N=5, 3, 3 and 3 hearts for IR, IR+CC, IR+Ex4 and IR+CC+Ex4 condition respectively) and the mean values per heart were used for the analysis in panel (c). (**c**) Mean distance from cell membrane to peak fluorescence intensity of mean Kir6.1 signal (also shown as red arrows in left plots). Scale bar is 10 µm.

### Signalling mediating remote ischaemic preconditioning

A signalling pathway consistent with the data presented above is schematised in Fig. 6. Ischaemia *in vivo* or OGD *ex vivo* inhibits Ca^2+^ pumping out of pericytes, activating pericyte contraction and capillary constriction. Ischaemic preconditioning releases GLP-1 from the gut^13–15^, which activates GLP-1 receptors on pericytes, raising the concentration of cAMP^34^ or activating PI3 kinase in the pericytes^15,19,35^ and thus activating K^+^-ATP channels (provided AMPK activation^24^ has inserted them into the surface membrane). K^+^ efflux then hyperpolarises the pericyte membrane, inhibiting Ca^2+^ influx through voltage-gated Ca^2+^ channels and thus relaxing pericytes.

**Figure 6.**
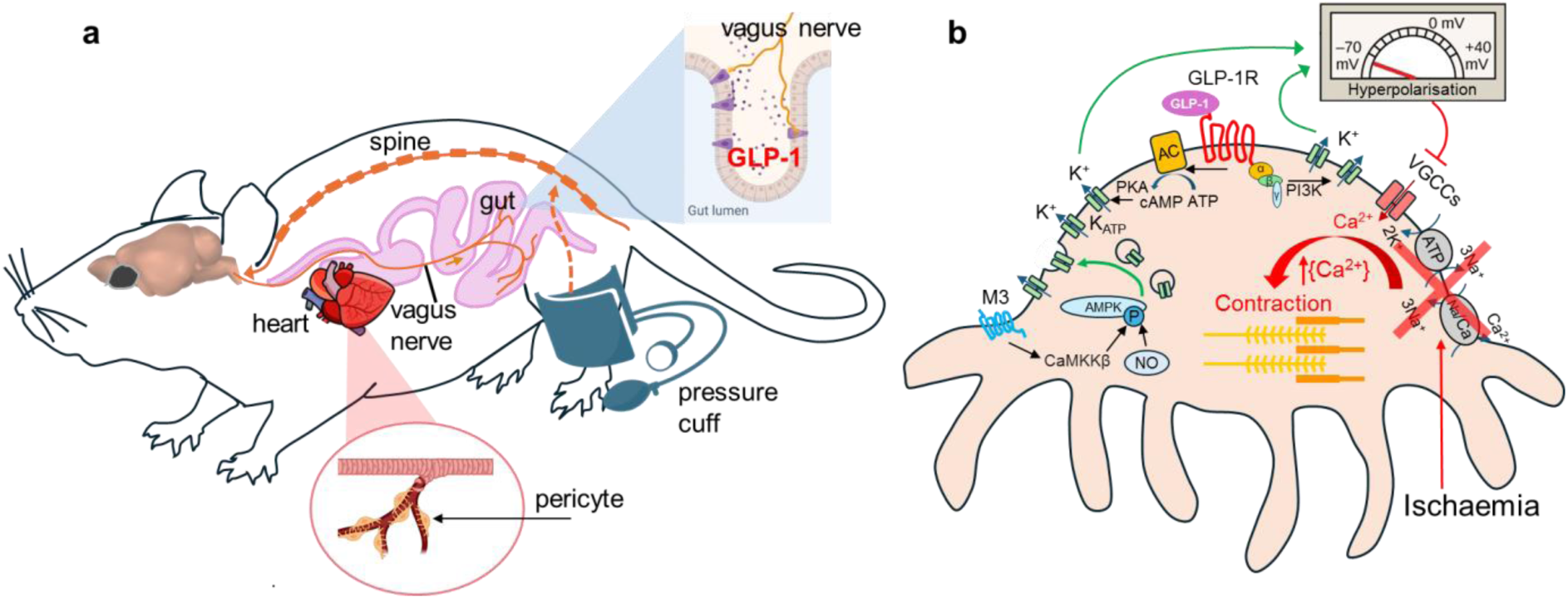
Signalling pathways characterised in this paper. (**a**) Schematic representation of brain-gut-heart neuro-humoral pathway in RPc cardioprotection. Limb ischaemia activates the RPc reflex via the spine, processed at the level of the brainstem. The neural stimulus is then relayed via the vagus to the gut, where GLP-1 is produced by enteroendocrine L-cells (inset: L-cells – purple; vagus –orange/brown). GLP-1 enters the systemic circulation with the blood/lymph and reaches the heart where it acts on the microvasculature to relax pericytes. (**b**) Molecular mechanisms downstream of GLP-1R activation on cardiac pericytes. Ischaemia *in vivo* or OGD *ex vivo* depletes intracellular ATP and inhibits Ca^2+^ pumping out of pericytes, activating pericyte contraction and capillary constriction. Ischaemic preconditioning releases GLP-1 from the gut, which activates GLP-1 receptors on pericytes, raising the concentration of cAMP or activating PI3 kinase, and thus activating K^+^-ATP channels (provided AMPK activity has inserted them into the surface membrane). K^+^ efflux then hyperpolarises the pericyte membrane, inhibiting Ca^2+^ influx through voltage-gated Ca^2+^ channels and thus relaxing pericytes. The activity of AMPK depends on muscarinic (M3) and nitric oxide mechanisms.

## Discussion

Following coronary ischaemia, lack of re-flow in the microcirculation after an upstream artery is unblocked has a negative impact on clinical outcome. In this study we have confirmed our earlier discovery that no-reflow after coronary ischaemia reflects pericyte-mediated constriction of capillaries^8,12^ (Fig. 2). This constriction will reduce blood flow by increasing the vascular resistance (both according to Poiseuille’s law and because the viscosity of the blood increases at small vessel diameters due to interactions of the cells suspended in the blood with the capillary wall^36^). In addition, it is likely that, as in the brain, there is an extra suppressive effect on blood flow by neutrophils and other cells stalling in capillaries near pericyte-induced constrictions^37,38^.

Remote ischaemic preconditioning (RPc) is an inter-organ protection phenomenon whereby ischaemia of an organ distant from the heart protects the myocardium against ischaemia/reperfusion injury. This phenomenon involves activation of parasympathetic vagal outflow, and release from the gut^14^ of GLP-1^15^ which, via the systemic circulation, reaches the heart and conveys cardioprotection. Cardioprotective factors have also been suggested to be released from the spleen under vagal control^39^, but GLP-1 is not produced in the spleen^40,41^, and a role for the spleen is controversial^42^. In rats activating GLP-1 receptors with Ex4 during reperfusion after coronary ischaemia reduces infarct size^43^, and in human patients giving GLP-1 after myocardial infarction or in heart failure improves outcome^44,45^, although the mechanism by which GLP-1 conveys cardioprotection has remained unclear^46^. Here, we demonstrate that GLP-1R mediated cardioprotection is at least partly mediated by its relaxing effect on pericytes on coronary capillaries. This is analogous to GLP-1R mediated relaxation of the coronary arteries^18^ and aorta^34^.

The long-lasting ischaemia-evoked constriction of capillaries by pericytes, and hence no-reflow, was reduced by RPc (Fig. 2g), which evokes GLP-1 release^15^. GLP-1 (or its analogue Ex4) is presumed to activate pericyte GLP-1 receptors and, by altering the concentration of cyclic AMP or by activating PI3 kinase^15,19,34,35^, thus activates ATP-gated K^+^ channels (Fig. 3) as in pancreatic beta cells^19^ (Fig. 6). As a result, K^+^ efflux hyperpolarises the cell, decreasing activation of voltage-gated Ca^2+^ channels, relaxing pericytes, and thus dilating capillaries near the pericyte somata where the majority of pericytes’ circumferential processes are located^47^. Involvement of ATP-gated K^+^ channels was shown both with pharmacological blockers and with pericyte-specific knockout of these channels. A similar restoration of capillary diameter towards normal size is seen in the short term when administering adenosine after coronary ischaemia^8^, probably also reflecting activation of ATP-gated K^+^ channels^48^, and administering adenosine or the ATP-gated K^+^ channel opener nicorandil improves outcome in human percutaneous coronary angioplasty by preventing no-reflow and thus improving ventricular function^49–51^.

Consistent with our LUXendin555 localisation data (Fig. 1), we assume for simplicity that the GLP-1 receptors mediating RPc are at least in part on the pericytes themselves, in order for them to adjust capillary diameter by altering pericyte contractile tone. The dilation of coronary capillaries by Exendin-4 in OGD is consistent with this GLP-1R agonist increasing coronary blood flow in humans^52^. It has previously been suggested that GLP-1 receptors are present on cardiac myocytes^53^, but deleting any such receptors did not abolish the protective effect of Exendin-4 on coronary ischaemia^53^ (which we attribute to increased coronary blood flow: Fig. 3). It has also been suggested that GLP-1Rs are located on endothelial cells of the endocardium^54^. While our imaging data (which were on the outer third of the ventricular myocardium where capillaries run more parallel to the section surface, making it easier to identify pericytes) suggested the presence of GLP-1 receptors on both pericytes and endothelial cells (Fig. 1), in any case gap junctional connections between endothelial cells and pericytes^55^ could conceivably allow activation of GLP-1Rs on endothelial cells to evoke activation of ATP-gated K^+^ channels in pericytes and thus to produce capillary dilation.

In order for activation of ATP-gated K^+^ channels to hyperpolarise the cell and thus relax pericytes, these channels need to be present in the surface membrane. It is unclear whether these channels are in the surface membrane normally, or whether they are inserted into the membrane only in conditions of ischaemia when AMP kinase activation is increased^24^. Blocking AMPK was found to remove the capillary-dilating protective effect of GLP-1R activation in OGD (Fig. 4a), probably because OGD-evoked AMPK activity^24^ is needed to insert ATP-gated K^+^ channels into the membrane^25,26^, and block of AMPK is known to prevent the reduction of infarct size by RPc^28^. Consistent with the protective effect of AMPK, knock-out or pharmacological block of the G-protein coupled receptor GPR39 was previously found to promote reflow and cardiac protection after ischaemia^56^. GPR39 inhibits cardiac AMPK^57^, so knock-out or block of GPR39 is expected to increase the surface membrane density of K_ATP_ channels in pericytes. In accord with this, with GPR39 not functioning, the diameter of capillaries was larger near pericyte somata^56^. Similarly, both NO and ACh muscarinic receptors are known to activate AMPK^30–33^ and, although blockers of NO synthase or of muscarinic receptors did not affect the capillary constriction induced by oxygen-glucose deprivation (Fig. 4b-c), they inhibited the capillary dilation evoked by GLP-1 receptor activation, consistent with these blockers preventing cardioprotection by RPc^15,29^. The effect we observed of the muscarinic blocker atropine, inhibiting the effect of GLP-1 receptor activation (Fig. 4c), in an isolated piece of ventricle, implies that there is tonic local release of ACh in the isolated heart, possibly via a non-neuronal cholinergic system in the ischaemic heart^58,59^. Consequently, damage-promoting effects of atropine in the whole animal^15^ may reflect actions intrinsic to the heart as well as suppression of vagally-induced GLP-1 release from the gut.

The increase in coronary blood flow that we suggest here is produced by GLP-1 has similarities to the increase in cerebral blood flow suggested to result from GLP-1 release during remote ischaemic preconditioning evoked before stroke^60^. Nevertheless GLP-1 analogues, while improving blood flow in pathology-affected tissues, may also have side effects, for example on the pancreas and gastrointestinal tract. Indeed, even in the heart, GLP-1R activation has been suggested to have actions other than opposing the effects of ischaemia-evoked blood flow reduction^61^. An increasing number of GLP-1 analogues are now being used in clinical practice, for disorders ranging from type 2 diabetes to obesity management and Alzheimer’s disease. It would be desirable to develop a version of these drugs that specifically target pericyte-mediated reduction of coronary blood flow, for use in cases of coronary ischaemia.

## Funding

This study was supported by a BHF Intermediate Basic Science Research Fellowship (FS/IBSRF/21/25060) to SM, and an ERC Advanced Investigator Award (740427, BrainEnergy), Wellcome Trust Senior Investigator Award (099222/Z/12/Z), a Rosetrees Trust grant on cardiac and renal ischaemia (M153-F2), and a BHF/UK-DRI Centre for Vascular Dementia Research grant to DA.

## Acknowledgements

We thank Frank Kirchhoff for NG2-Cre mice, Akiko Nishiyama and Dirk Dietrich for NG2-dsRed mice, Andrew Tinker for Kir6.1 (flx/flx) mice, Elisa Avolio for providing human pericyte culture, David Hodson for providing Luxendin555 probe, Thomas Kampourakis for suggesting the use of Mavacamten, Stuart Martin for genotyping and Wolfgang Langhans for advice during this work.

## Data Availability

All relevant data generated in this study are available within the article and its Supplementary Information files. The source data are provided with this paper.

## Methods

Experiments followed the guidelines of the European Commission Directive 2010/63/EU (European Convention for the Protection of Vertebrate Animals used for Experimental and Other Scientific Purposes) and the UK Home Office (Scientific Procedures) Act (1986), with approval from the Institutional Animal Care Committee.

### Animal preparation

Adult male Sprague-Dawley rats weighing 220-250g or transgenic mice (see below) were anaesthetized with pentobarbital sodium (induction 60 mg/kg i.p. for rats, 90 mg/kg i.p. for mice; maintenance 10-15 mg/kg/h i.v.). The right carotid artery and jugular vein were cannulated for blood pressure (BP) measurement and drug administration, respectively. Stable blood pressure and heart rate were maintained, and level of anaesthesia was monitored by paw pinch response. The trachea was cannulated, and animals were mechanically ventilated with room air using a positive pressure ventilator (Harvard rodent ventilator, tidal volume of 1 ml/100 g of body weight, ventilator frequency ∼60 strokes min^−1^ for rats, ∼150 strokes min^−1^ for mice). Arterial BP and ECG were recorded using a Power1401 interface and *Spike2* software (Cambridge Electronic Design), and body temperature was maintained at 37.0±0.5°C. The heart was exposed via a left thoracotomy and a 4-0 (rats) or 8-0 (mice) prolene suture was passed around the left anterior descending (LAD) coronary artery to induce a temporary occlusion. Rats were subjected to 45 mins of LAD ligation followed by 15 mins of reperfusion to study no-reflow phenomena. To determine the effects of Exendin-4 (a GLP-1R agonist) and Compound C (an AMPK inhibitor) on the localisation of K_ATP_ channels in ischaemia, mice were subjected to 40 mins of LAD ligation followed by 10 mins of reperfusion, with Compound C (Dorsomorphin 2HCl, APExBIO; 10 mg/kg, i.p., administered 1 h before LAD ligation) and/or Exendin-4 (Tocris; 5 µg/kg i.p., given 30 mins before LAD ligation). To determine the extent of myocardial ischaemia/reperfusion damage, mice were subjected to 30 mins of LAD ligation followed by 90 mins reperfusion. Successful LAD occlusion was confirmed by paling of the myocardial tissue distal to the suture, elevation of the ST-segment in the ECG, and an immediate 15–30 mm Hg fall in the BP. To establish remote ischaemic preconditioning, blood supply to the lower limbs was interrupted for 15 mins by placing clamps on both femoral arteries at the proximal level ∼1 cm below the inguinal ligament 25 mins before the onset of coronary ischaemia. The sham-RPc procedure involved dissection of both femoral arteries without occlusion. GLP-1R antagonist Exendin(9–39) (Tocris, Ex(9-39), 50 µg/kg, i.v.) or vehicle control was administered 40 mins before the onset of coronary ischaemia. Control (sham operated) animals underwent the same procedures, except that after the suture was passed around the LAD coronary artery it was not drawn tight to occlude the vessel. Ischaemia and sham animals were alternately interleaved.

### Infarct size measurement

In mice, at the end of reperfusion period, the LAD was re-ligated, and 5% Evans Blue dye solution (100 µl) was infused via jugular vein to determine area at risk. The animal was then given an anaesthetic overdose (pentobarbital, 200 mg^-1^ kg^-1^, i.v.), the heart was excised, briefly frozen and sectioned into 5-6 transverse slices from apex to base. The area at risk was demarcated by the absence of Evans Blue staining. Heart slices were then incubated with 1% 2,3,5-triphenyltetrazolium chloride (TTC) in Tris buffer (pH 7.4) for 15 mins at 37 °C and fixed in 4% formalin for 24 h. Viable myocardium is stained red by TTC, whereas necrotic myocardium appears white. The area at risk and the necrotic area were determined by computerized planimetry, normalized to the weight of each slice, with the degree of necrosis (i.e. infarct size) expressed as the percentage of area at risk, as described^13,14^.

### Animal perfusion and tissue processing for imaging

After the ischaemia/reperfusion procedure, rats were given an overdose of pentobarbital sodium and transcardially perfused with 200 ml saline followed by 200 ml 4% paraformaldehyde (PFA) for fixation and then by a solution of 5% gelatine (Sigma-Aldrich, G2625) containing 0.25% FITC-albumin conjugate (Sigma-Aldrich, A9771) for visualization of the perfused coronary microvasculature. The initial perfusion with calcium-free saline ensures that the heart is stopped in diastole, which is also evident from the large volume of left ventricle visible at cross-section. The hearts were then fixed overnight in 4% PFA, and 150 µm transverse sections were obtained for immunofluorescence staining. The labeling was done using anti-NG2 antibodies (Merck Millipore, AB5320, 1:200) for pericytes and isolectin B_4_-Alexa Fluor 647 (Molecular Probes, I32450, 1:200) for the capillary basement membrane.

Mice used in Kir6.1 membrane localisation study were given an anaesthetic overdose at the end of 10 mins reperfusion and transcardially perfused with 20 ml saline followed by 5 ml 4% PFA for fixation, the hearts were fixed overnight in 4% PFA and sliced into 150 µm transverse sections. The labeling was done using anti-PDGFRb antibodies (R&D Systems, AF1042, 1:50) for pericytes, isolectin GS-IB4-biotinylated (Life Technologies, 121414, 1:200) for capillary basement membrane and anti-Kir6.1 (Alomone Labs, APC-105, 1:100) to localise the K_ATP_ channels, following heat-induced epitope retrieval (using 10 mM sodium citrate buffer at pH 6.0 for 20 mins at 80°C).

To visualise GLP-1Rs in mouse pericytes, wild type mice were injected with Luxendin555 probe subcutaneously^62^ (100 pmol/g body weight) and perfused-fixed (as described above) 2 h after Luxendin555 administration. Tissue was processed as described above; the pericytes and blood vessels were labelled using anti-NG2 antibodies (Abcam, ab275024, 1:200) and isolectin B_4_-Alexa Fluor 488 (Molecular Probes, I21411, 1:200) respectively (Fig. 1a). In a separate set of experiments, heart tissue from NG2-dsRed mice was processed as described above and stained with anti-GLP-1R antibody (Abcam, ab218532, 1:200) following heat-induced epitope retrieval as above.

To visualise GLP-1Rs in human pericytes, pericytes isolated from cardiac surgery waste tissue (cell isolation methods and characterisation as described previously^63^) were cultured to semi-confluency, fixed with 4% PFA for 20 mins and labelled with the anti-human PDGFRβ antibody (Santa Cruz, sc-374573, 1:50) and Fluorescent Exendin-4 (Fluorescein-Trp25-Exendin-4, FLEX; Eurogentec, AS-63899, 100 nM).

### Imaging vessels after *in vivo* ischaemia experiments

Images of the left ventricular capillary bed in rats were acquired using laser scanning confocal microscopy (Zeiss LSM 700) and analysed using FIJI software (ImageJ 1.53c, NIH). To quantify blood volume across the left ventricular myocardium, low power z-stack images were taken of an entire transverse section of each heart (using a 1X air objective), maximum intensity projected.

For quantification of the percentage of capillaries that were perfused, three randomly selected regions of the outer third of the myocardium of the anterior left ventricular wall were imaged, as this region of myocardium consistently included the ischaemic area (which showed visible pallor and oedema of the myocardium). The person quantifying the images was blinded to the condition that the heart was exposed to. Blockages of flow in the ischaemic area at risk were identified by abrupt terminations in FITC-albumin signal (Fig. 2a, b).

### Transgenic mice

Some experiments used NG2-dsRed mice^64^, kindly provided by Akiko Nishiyama (University of Connecticut) via Dirk Dietrich (University of Cologne), in which pericytes fluoresce red, facilitating their identification. Pericyte-specific Kir6.1 knockout mice were generated by crossing tamoxifen-inducible NG2-Cre^ERT2^ knock in mice^65^ (kindly donated by Frank Kirchhoff, University of the Saarland) and floxed (flx) Kir6.1 mice (kindly donated by Andrew Tinker, QMUL) to allow deletion of Kir6.1 in NG2-expressing pericytes after oral gavage of tamoxifen. Tamoxifen dissolved in corn oil was given at 100 mg/kg body weight by oral gavage once per day for three consecutive days to adult >P21 mice. Experiments were performed from 2 weeks after tamoxifen administration.

Genotypes were identified by polymerase chain reaction on genomic DNA from ear snips of NG2-cre(+/-)/Kir6.1(flx/flx), and littermate control animals NG2-cre(+/-)/Kir6.1(-/-) using the following primers: for NG2-Cre, WT sense AGAGATCCTGTCCACAGAGTTCT and antisense GCTGGAGCTGACAGCGGGTG and generic Cre, sense CACCCTGTTACGTATAGCCG and antisense GAGTCATCCTTAGCGCCGTA (NG2 WT band 557 b.p., Cre band 330 b.p.); for Kir6.1, sense 5’- GAGTCTTAACTCAGTTCTGGAGGACCAACA-3’ and antisense 5’-AGCGAAGAAAACTGCTTCCTGTTCATTAAG-3’ (Kir6.1 flx band 600 bp, WT band 474 bp).

### Live *ex vivo* tissue imaging

Adult mice (P100-P120) of both sexes were used in the study. The heart of a freshly euthanised mouse was surgically removed and placed into ice-cold Tyrode’s solution containing the following components (mM): 142.5 NaCl, 2.5 KCl, 0.5 MgCl_2_, 0.33 NaH_2_PO_4_, 5 glucose, 2 Na lactate, 0.1 Na pyruvate, 10 Hepes and 1.8 CaCl_2_ (pH set to 7.4 with NaOH). After being cleaned of connective tissues, the heart was transferred into a Sylgard-coated prechilled (4°C) chamber containing Tyrode’s solution and pinned down onto the Sylgard layer of the chamber. The right ventricular wall was dissected, and the right coronary artery exposed and cannulated using a fire-polished bent glass pipette prefilled with Tyrode’s solution. The right ventricular wall was used for imaging as it was thinner than the left ventricular wall, which made it easier to flatten under the objective and ensured full immersion of the tissue in the bath superfusion solution. The pipette used to cannulate the artery was then connected to a fluid-filled line, and intralumenal pressure was created by elevating a cylinder connected to the line with Tyrode’s solution to a height equivalent to 80 mm Hg (109 cm H_2_O) (Fig. 3a). The vascular lumen perfusate was Tyrode’s solution (composition as above). The bath superfusate contained a physiological saline solution (PSS; containing (mM) 114.5 NaCl, 2.5 KCl, 1.2 MgSO_4_, 1.2 NaH_2_PO_4_, 24 NaHCO_3_, 10 glucose, 1 Na lactate and 1.8 CaCl_2_) was maintained at 32-34°C with a heater and temperature controller and bubbled with 5% CO_2_, 20% O_2_ and 75% N_2_ to maintain a pH of 7.4. The bath solution was perfused at ∼5 ml/min. Experimental measurements were started after ∼30-60 mins stabilization. To visualise the vessels, Alexa Fluor 488-conjugated WGA (Thermo Fisher Scientific, W11261; 20 µg/mL, 30 mins) was included in the luminal buffer during the stabilisation period. For imaging of the tissue from animals not expressing dsRed in pericytes, an additional step of 20 mins incubation with isolectin B_4_-Alexa Fluor 594 (Molecular Probes, I21413, 10 µg/ml in superfusate solution) was employed. To block cardiac myocyte contraction, a modulator of cardiac myosin, Mavacamten^66^ (MYK-461, 10µM, MedChemExpress), was added to the luminal solution.

Two-photon excitation used a Newport-Spectra Physics Mai Tai Ti:Sapphire Laser pulsing at 80 MHz, and a (Zeiss LSM780, Oberkochen, Germany) microscope with a 20× water immersion objective (NA 1.0). Fluorescence was excited using 920 nm wavelength for DsRed, and 820 nm for Alexa Fluor 488. Mean laser power under the objective was <35 mW. Z-stack images were acquired every 2 mins. Baseline for each condition was 5 mins. The *ex vivo* model of tissue ischaemia was induced by oxygen/glucose deprivation (OGD) – the tissue was exposed to perfusate solution saturated with 95% N_2_/5% CO_2_ gas mixture containing 10 mM sucrose instead of glucose. The period of OGD lasted for 25 mins, after which GLP-1R agonist was added to the perfusate solution, and OGD continued for another 25 mins. Exendin-4 (Ex4; 100 nM, Tocris) was used as GLP-1R agonist. *N*(gamma)-nitro-L-arginine methyl ester (L-NAME; 100 μM, Sigma) was used to inhibit nitric oxide synthesis. Atropine sulphate (1 µM, Sigma) was used to block muscarinic receptors. Compound C was used to inhibit AMPK (5 µM, APExBIO). K_ATP_ channels were activated with pinacidil (100 µM, AOBIOUS) and blocked with glibenclamide (20 µM, Santa Cruz). Endothelin-1 (20 nM) was applied either through the vascular lumen or in the superfusate, and the constriction evoked did not differ significantly (22.3±7.3% constriction when ET-1 was applied in the lumen solution compared to 24±5.4% constriction when ET-1 was applied in the bath solution). For mechanistic experiments, ET-1 and pinacidil were applied via the capillary lumen, while all other drugs were applied in the superfusion solution.

### Image analysis

For low power blood volume analysis following *in vivo* experiments, 12 regions of interest (ROI) were drawn clockwise around the left ventricle (when looked at from above) from the mid-point of the septum^8^ (as in Fig. 2a, d), and the mean intensity of FITC-albumin signal was recorded for each ROI, and normalised to the highest intensity measured in any ROI. These data were averaged over hearts and renormalized so that the mean value in positions 1–3 of Fig. 2c was 1. This signal is assumed to be proportional to the volume of blood perfusing the myocardium. To compare perfusion in the ischaemic zone, we averaged the plotted values over ROIs 7–10 for each heart in the different conditions, and compared the mean values averaged over all the hearts studied in each condition.

Pericytes were identified by their morphology (spatially isolated cells located outside capillaries) as confirmed by IB_4_ labeling, or by antibody labeling for their characteristic markers NG2 or PDGFRβ, or by expression of dsRed under the NG2 promoter. To quantify the percentage of perfused capillaries, we counted the number of filled and unfilled vessels crossing a line drawn through the centre of each image perpendicular to the main capillary axis (as in Fig. 2b, e).

Capillary diameters and Kir6.1 immunofluorescence intensity profiles were analysed using FIJI software (ImageJ 1.53c, NIH). Vessel diameter was defined using a line drawn across the vessel as the width of the intraluminal dye fluorescence (WGA-Alexa Fluor 488) at pericyte somata. To assess the Kir6.1 immunofluorescence intensity profiles, PDGFRβ immunostaining was used as a reference point to localise the cell surface. 10 regions of interest (ROI) per pericyte were drawn as lines in the PDGFRβ channel starting from the cell surface and directed towards the centre of the cell, the ROIs were then transferred to the Kir6.1 channel and the fluorescence intensity profile of each line was measured and averaged over all the lines per pericyte. The values of the resulting distance from the cell membrane to the peak Kir6.1 immunofluorescence intensity were averaged for each heart were then used for the analysis (Fig. 5).

### Statistics

Statistical analysis was performed using GraphPad Prism 9 software. Data normality was assessed with Shapiro-Wilk tests. Comparisons of normally distributed data were made using 2-tailed Student’s t-tests. Equality of variance was assessed with an F test, and heteroscedastic t-tests were used if needed. Data that were not normally distributed were analysed with Mann-Whitney tests. P values were corrected for multiple comparisons using the Šidák method or a procedure equivalent to the Holm-Bonferroni method (for N comparisons, the most significant p value is multiplied by N, the 2nd most significant by N-1, the 3rd most significant by N-2, etc.; corrected p values are significant if they are less than 0.05). Numbers of animals are denoted as N, and numbers of pericytes or capillaries as n. In all statistical tests, animals (N) were the statistical unit. Data are presented as mean values ± SEM.

## Notes

### Competing Interest Statement

The authors have declared no competing interest.

